# An updated resource of 180K soybean SNP genotyping array based on the T2T reference genome

**DOI:** 10.1101/2025.07.29.667553

**Authors:** Ji-Hun Hwang, Sungwoo Lee, Ju Seok Lee, Kyung Do Kim

## Abstract

Single nucleotide polymorphism (SNP) genotyping has revolutionized crop improvement by enabling high-resolution genomic analyses and accelerating breeding programs. In soybean (*Glycine max* (L.) Merr.), a globally important legume crop, existing genotyping data for 180,961 SNP markers from the Korean soybean core collection were generated using the outdated Williams 82 reference genome version 1 (Wm82.v1), which contains numerous assembly gaps and misassemblies that limit genomic resolution. While high-quality reference genomes including Wm82.v4 and Wm82.v6 (telomere-to-telomere assembly) are now available, the valuable existing SNP array data have not been integrated with these improved genomic resources. Here we show successful remapping of the 180K SNP array data to both Wm82.v4 and Wm82.v6 reference genomes through sequence-based alignment of flanking regions. We extracted flanking sequences from SNP marker positions in Wm82.v1 and mapped them to the newer reference versions based on sequence similarity, excluding markers with mapping failures, allele mismatches, low-identity alignments, or multiple mappings, which resulted in successful mapping of 175,202 and 175,763 markers to Wm82.v4 and Wm82.v6, respectively. We also remapped genotype data from 927 soybean accessions (497 USDA-GRIN accessions and 430 Korean core collection accessions) to both reference versions. This updated SNP dataset provides the soybean research community with a comprehensive genomic resource that leverages both existing genotyping investments and state-of-the-art reference genome assemblies for enhanced crop improvement and genomic studies.

## Introduction

Soybean (*Glycine max* (L.) Merr.), a major crop worldwide, is an essential source of protein and oil for human nutrition, animal feed, and industrial applications. Advances in genomic technologies have substantially improved our understanding of soybean genetics, aiding in the identification of traits linked to yield, stress resistance, and quality improvement [1–2]. Among these technologies, single nucleotide polymorphism (SNP) arrays have enabled high-throughput genotyping and trait mapping in soybean breeding programs [3].

The 180K SNP array [4], developed using the Williams 82 (Wm82) genome version 1 (Wm82.v1) and version 2 (Wm82.v2) [5], is a pivotal resource for genetic studies in the Korean soybean core collection [6]. However, the Wm82.v1 and Wm82.v2 genomes, published in the early stages of soybean genomics, are characterized by incomplete assemblies and structural errors that limit the resolution and accuracy of downstream analyses. These challenges are compounded by the palaeopolyploid nature of the soybean genome, which includes extensive duplications and structural rearrangements from past whole-genome duplications. The use of short-read sequencing technologies further constrained the ability of the assembly to resolve repetitive and complex regions, leaving significant gaps and misassemblies.

The evolution of sequencing technologies, particularly the advent of long-read and high-fidelity sequencing, has dramatically improved the quality of genome assemblies [7–8]. Subsequent releases of the Wm82 genome, such as Williams 82 version 4 (Wm82.v4) and version 6 (Wm82.v6), represent significant milestones, with Wm82.v6 achieving a telomere-to-telomere (T2T) level of completeness [9–10]. These updated reference genomes resolved many structural issues inherent to Wm82.v1, thus providing a more robust framework for genomic analysis. Despite the availability of high-quality genome assemblies, many legacy SNP datasets remain anchored to outdated references. This disconnect hinders their integration with modern genomic tools.

Thus, this study aimed to reposition the 180K SNP array dataset, which was originally genotyped on older versions of the Wm82 genome, onto the latest versions 4 and 6. By performing this liftover process, we sought to improve the accuracy of SNP positions, resolve inconsistencies arising from genome version differences, and ensure compatibility with modern genomic analyses. Repositioning of these datasets is expected to enhance the reliability of marker-trait associations, optimize genomic selection strategies in breeding programs, and facilitate functional genomic studies by aligning genotypic data with the most up-to-date reference genomes.

## Materials and methods

### Data collection

Affymetrix Axiom® 180K SoyaSNP array data were collected from the previous study [4]. The genotype data of 4,234 soybean accessions, derived from the Affymetrix Axiom® 180K SoyaSNP array, were collected from the previous study [6]. The genotype data of 430 Korean soybean core collection and 497 plant introduction (PI) soybean collection were obtained from the dataset of 4,234 soybean accessions. Various versions of the Wm82 genome and genes were collected from SoyBase (www.soybase.org) [11]. Transposable element (TE) sequences [12] of soybean were also collected from SoyBase.

### 180K Axiom® SoyaSNP array liftover

To liftover the 180K SoyaSNP array data to each Wm82 genome, we extracted the 50 bp flanked upstream and downstream sequences at the SNP position of Wm82.v1 using bedtools [13]. BLAT was used to identify homologous regions between the flanked SNPs from the assembly version Wm82.v1 and the other assembly versions of Wm82 [14]. The UCSC axtChain was used to chain the alignment blocks and identify the collinearity between genomes [15]. Of the 180K SNP data, 170K SNPs representing genetic variation within the Korean soybean core collection in VCF format were lifted using the Picard tool, LiftoverVcf (v2.22.8), and chain information (http://picard.sourceforge.net/). The remaining SNPs of the 180K data were lifted based on positional information from the chain file.

### Liftover validation

The flanking sequences of SNPs redundantly aligned to multiple locations between Wm82.v4 and Wm82.v6 were filtered. The lifted SNPs were validated using an in-house script that compared the flanked sequences from the source and target assembly using SAMtools [16]. After the validation, flanking sequences of SNPs that show identity less than 100% between Wm82.v1 and the target Wm82 genome sequence were filtered. The density of the genes and lifted markers at Wm82.v6 were visualized using the RIdeogram package in R [17]. Bedtools was used to categorize lifted markers as genic, near-genic, or intergenic at Wm82.v4 and Wm82.v6. Gene synteny between Wm82.v4 and Wm82.v6 was identified using MCScan [18]. The NUCmer package of MUMmer was employed to align the intergenic regions between Wm82.v4 and Wm82.v6 [19]. Syri was used to identify genomic rearrangements and collinear regions between genomes [20]. JBrowse2 was employed to visualize the microstructure of each genome [21]. RepeatMasker was used to identify the location of TEs [22]. NABIS 2, a high-performance computer from the Rural Development Administration of South Korea, was utilized for the computational analysis.

### Statistical analysis

Minor allele frequencies (MAFs) for each marker position in the soybean collection’s genotype data were calculated using an in-house script.

## Results

### Liftover of 180K Axiom® SoyaSNP array

Of the 180,961 SNP markers from the Affymetrix Axiom® 180K SoyaSNP array genotyped onto Wm82.v1, 5,759 and 5,198 markers failed to be lifted to Wm82.v4 and Wm82.v6, respectively (Fig 1). Of these unmapped markers, 3,081 in Wm82.v4 and 2,974 in Wm82.v6 failed because of mapping to multiple genomic loci, and 1,728 and 1,267 were completely unmapped to the target genome. By contrast, 176,152 and 176,720 markers were initially lifted to Wm82.v4 and Wm82.v6, respectively. After the initial liftover, an additional 675 markers were filtered from both Wm82.v4 and Wm82.v6 owing to SNP allele divergence. In addition, 275 and 282 markers that did not exhibit 100% identity with the Wm82.v1 genome were filtered at Wm82.v4 and Wm82.v6. The 275 markers that failed to be lifted to Wm82.v4 showed sequence identities ranging from 50% to 70%. Meanwhile, 280 markers among the 282 markers that failed to be lifted to Wm82.v6 showed sequence identities ranging from 50% to 70%, and the 2 remaining markers had sequence identities ranging from 70% to 90% (S1 Table). The total number of retained markers did not significantly differ from the results obtained with a 100% identity threshold. Ultimately, 175,202 and 175,763 markers were retained in Wm82.v4 and Wm82.v6, respectively (Table 1).

**Fig 1.**
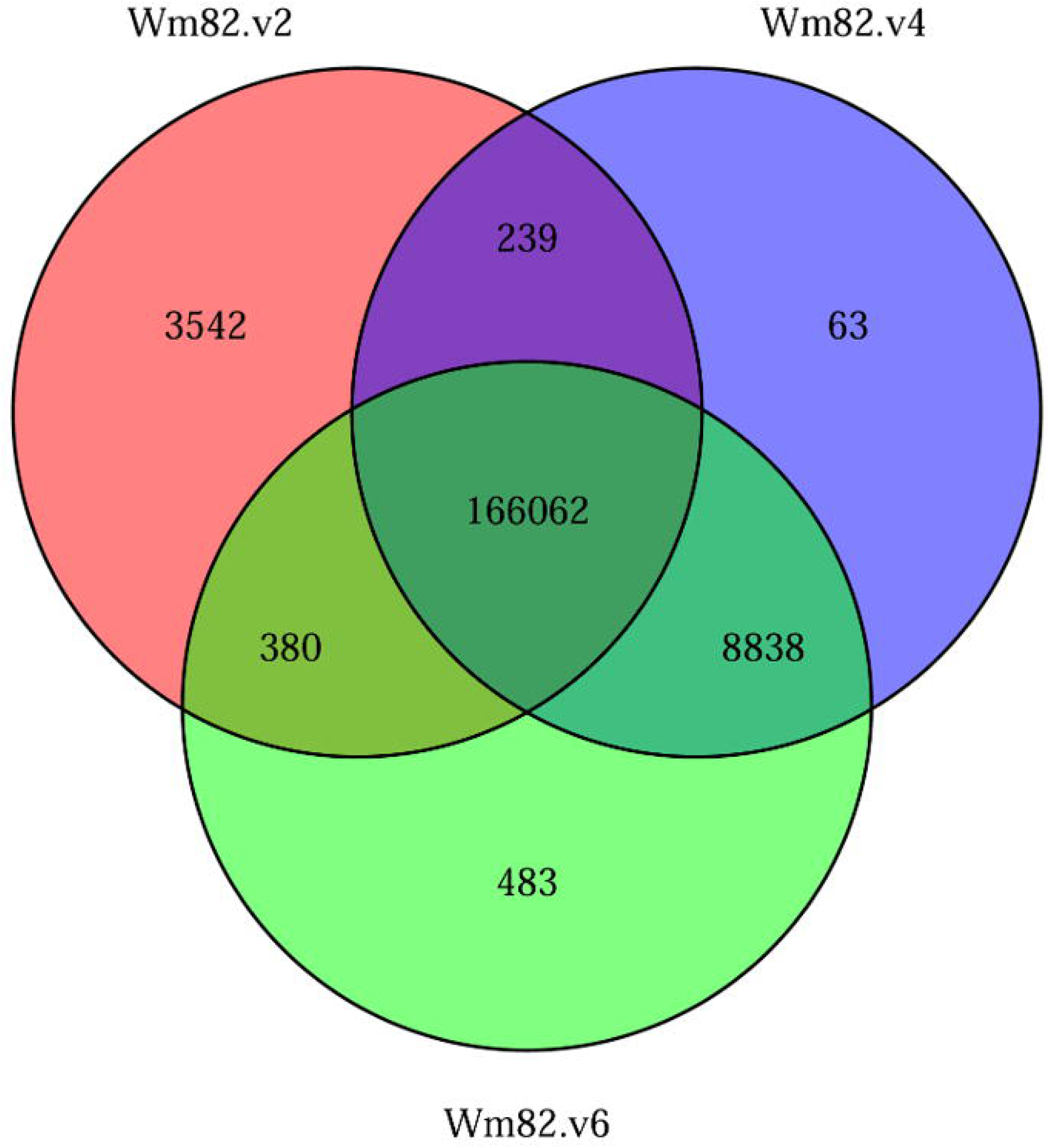
Overview of SNP markers with liftover failures.

**Table 1.** Summary of the 180K SNP array data liftover.

| Chromosome | Total SNPs |  | Lifted SNPs |  |  |  |  |  |
| --- | --- | --- | --- | --- | --- | --- | --- | --- |
|  | (Wm82.v1)<sup>ab</sup> |  | Wm82.v2<sup>ab</sup> |  | Wm82.v4 |  | Wm82.v6 |  |
| Chr01 | 8,935 | 4.94% | 8,379 | 4.63% | 8,797 | 4.86% | 8,833 | 4.88% |
| Chr02 | 10,224 | 5.65% | 9,514 | 5.26% | 9,857 | 5.45% | 9,885 | 5.46% |
| Chr03 | 8,417 | 4.65% | 7,673 | 4.24% | 7,737 | 4.28% | 7,772 | 4.29% |
| Chr04 | 8,638 | 4.77% | 8,399 | 4.64% | 8,567 | 4.73% | 8,597 | 4.75% |
| Chr05 | 8,024 | 4.43% | 7,577 | 4.19% | 7,653 | 4.23% | 7,727 | 4.27% |
| Chr06 | 9,906 | 5.47% | 9,392 | 5.19% | 9,607 | 5.31% | 9,693 | 5.36% |
| Chr07 | 8,588 | 4.75% | 8,126 | 4.49% | 8,209 | 4.54% | 8,234 | 4.55% |
| Chr08 | 10,996 | 6.08% | 10,475 | 5.79% | 10,712 | 5.92% | 10,777 | 5.96% |
| Chr09 | 8,996 | 4.97% | 8,621 | 4.76% | 8,912 | 4.92% | 8,962 | 4.95% |
| Chr10 | 9,309 | 5.14% | 8,820 | 4.87% | 9,099 | 5.03% | 9,153 | 5.06% |
| Chr11 | 8,412 | 4.65% | 7,481 | 4.13% | 8,172 | 4.52% | 8,222 | 4.54% |
| Chr12 | 7,708 | 4.26% | 7,222 | 3.99% | 7,495 | 4.14% | 7,520 | 4.16% |
| Chr13 | 10,885 | 6.02% | 10,338 | 5.71% | 10,595 | 5.85% | 10,687 | 5.91% |
| Chr14 | 7,817 | 4.32% | 7,195 | 3.98% | 7,560 | 4.18% | 7,604 | 4.20% |
| Chr15 | 10,136 | 5.60% | 9,423 | 5.21% | 9,835 | 5.43% | 9,864 | 5.45% |
| Chr16 | 7,590 | 4.19% | 6,980 | 3.86% | 7,348 | 4.06% | 7,367 | 4.07% |
| Chr17 | 8,906 | 4.92% | 8,347 | 4.61% | 8,663 | 4.79% | 8,682 | 4.80% |
| Chr18 | 9,957 | 5.50% | 9,103 | 5.03% | 9,518 | 5.26% | 9,576 | 5.29% |
| Chr19 | 8,719 | 4.82% | 8,166 | 4.51% | 8,480 | 4.69% | 8,503 | 4.70% |
| Chr20 | 8,212 | 4.54% | 7,797 | 4.31% | 8,072 | 4.46% | 8,105 | 4.48% |
| Scaffolds | 575 | 0.32% | 1,185 | 0.65% | 314 | 0.17% | 0 | 0.00% |
| **Total** | 180,950 | - | 170,213 | 94.07% | 175,202 | 96.82% | 175,763 | 97.13% |

<sup>a</sup>A data previously reported [4].
<sup>b</sup>Chloroplast markers were excluded.

The liftover rates of the markers in the Wm82.v4 and Wm82.v6 genomes were 96.82% and 97.13%, respectively. Among the lifted markers, 162,020 and 162,758 were lifted as forward strands, whereas 13,182 and 13,005 were lifted as reverse strands in Wm82.v4 and Wm82.v6, respectively (S2 Table). Among all the lifted markers, 166,062 were successfully mapped to Wm82.v2, Wm82.v4, and Wm82.v6 (Fig 2). Moreover, 9,457 markers were lifted to at least two different Wm82 versions. Specifically, 8,838 markers were lifted to both Wm82.v4 and Wm82.v6, 380 to Wm82.v2 and Wm82.v6, and 239 to Wm82.v2 and Wm82.v4. Additionally, 4,088 markers were uniquely lifted to a single Wm82 version, including 3,542 markers to Wm82.v2, 63 markers to Wm82.v4, and 483 markers to Wm82.v6. Furthermore, 1,354 markers could not be lifted to any of the Wm82.v2, Wm82.v4, and Wm82.v6 versions.

**Fig 2.**
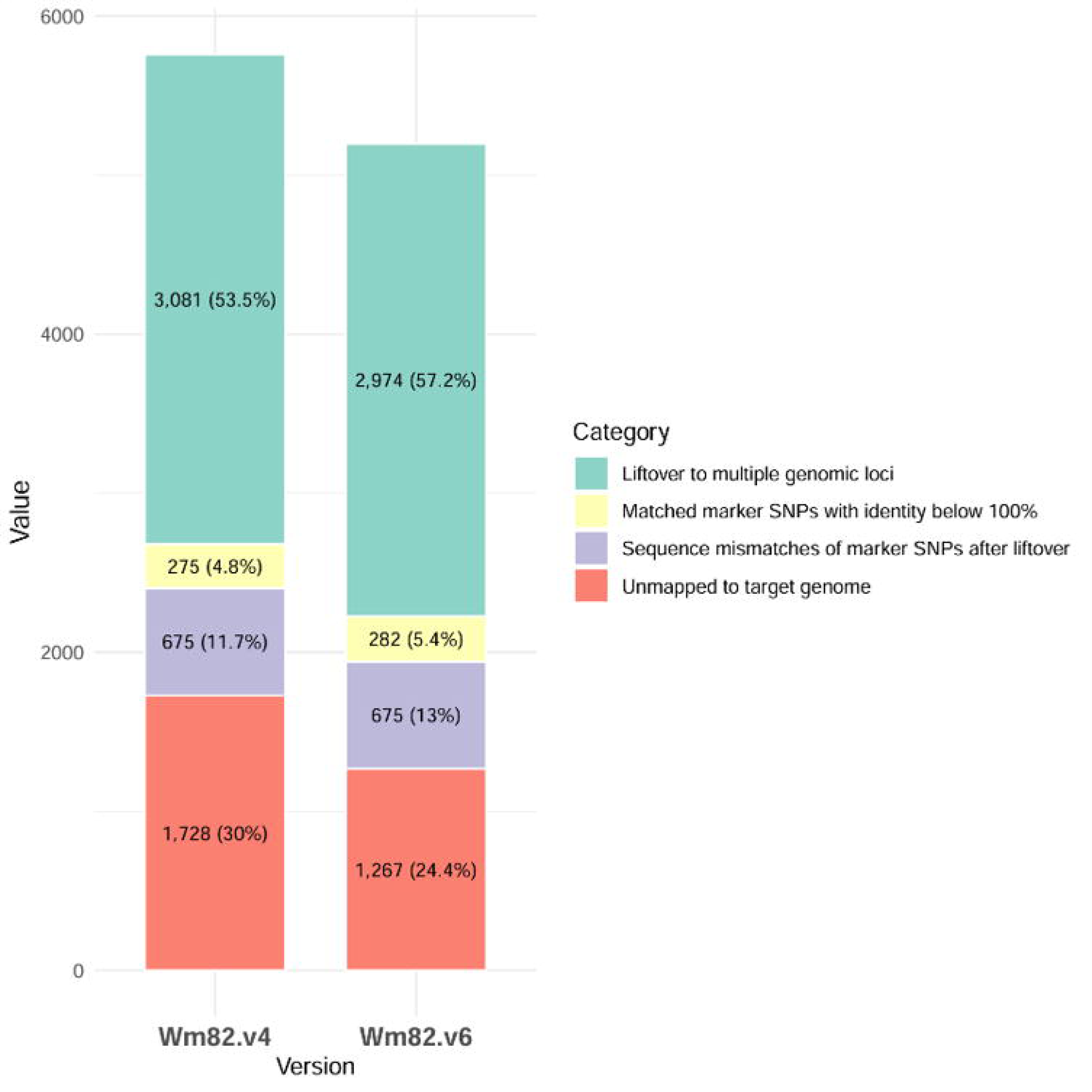
Venn diagram of the lifted SNP marker across the various versions of Wm82 genomes.

For chromosome-level comparison, we evaluated the number of markers retained on the same chromosome from Wm82.v1 to the target genome (S3 Table). Of the 8,935 markers genotyped on chromosome 1 of Wm82.v1 in a previous study, 97.88% and 98.27% of the markers were lifted to chromosome 1 in Wm82.v4 and Wm82.v6, which exhibited the highest success rates among the chromosomes. Of these, 8,746 and 8,780 markers were originally from chromosome 1, whereas 51 and 53 were lifted from the Wm82.v1 scaffolds. By contrast, chromosome 3 showed the lowest liftover rates of 91.85% and 92.17% for Wm82.v1 markers lifted to Wm82.v4 and Wm82.v6, respectively. The most significant difference in liftover rates between Wm82.v4 and Wm82.v6 was observed on chromosome 6, with rates of 96.64% and 97.70%, respectively, reflecting a 1.06% difference. Finally, we identified markers that were originally genotyped on scaffolds in Wm82.v1 but were lifted over to chromosomes in Wm82.v4 and Wm82.v6 (S4 Table). Of 575 markers of the Wm82.v1 scaffolds, 51 markers remained at the Wm82.v4 scaffolds, while 350 markers were lifted to the chromosome. However, in Wm82.v6, a T2T-level genome without any scaffold, 456 markers were lifted from the scaffolds to the chromosome. Among the originally genotyped markers on scaffolds in Wm82.v1, a total of 174 and 119 markers failed to be lifted to Wm82.v4 and Wm82.v6, respectively.

### Application of SNP liftover to the soybean collections

Among the 175,202 and 175,763 SNP markers that were lifted to Wm82.v4 and Wm82.v6, 166,301 and 166,442, respectively, were present among the 170,223 markers genotyped in the soybean collection from Wm82.v1. As a result, without duplicates, 166,681 markers were lifted to the markers of the soybean collection. In total, 166,062 markers were lifted to both Wm82.v4 and Wm82.v6, whereas 239 and 380 markers were lifted separately. For comparison, the lifted markers were classified based on allele frequency. We first examined the allele frequency in the 430 Korean soybean core collection (Table 2).

**Table 2.** Categorization of lifted markers to 430 Korean soybean core collection using MAF.

| SNP Type | MAF | Number of SNPs |  |  |  |
| --- | --- | --- | --- | --- | --- |
|  |  | Wm82.v4 |  | Wm82.v6 |  |
| Lifted<br>to 430 Korean<br>Soybean Core<br>Collection | ~0.01 | 62,998 | 37.88% | 63,063 | 37.89% |
|  | 0.01–0.05 | 22,990 | 13.82% | 22,995 | 13.82% |
|  | 0.05–0.10 | 17,123 | 10.30% | 17,148 | 10.30% |
|  | 0.10–0.20 | 24,328 | 14.63% | 24,359 | 14.64% |
|  | 0.20~ | 38,862 | 23.37% | 38,877 | 23.36% |
| **Total** |  | 166,301 | - | 166,442 | - |

A total of 62,998 and 63,063 markers from Wm82.v4 and Wm82.v6, respectively, which were lifted to the 430 Korean soybean core collection, exhibited MAFs below 1%. Additionally, 22,990 and 22,955 markers had MAFs between 1% and 5%; 17,123 and 17,148 within the 5%–10% range; and 24,328 and 24,359 between 10% and 20%. In total, 38,862 and 38,377 markers with MAFs greater than 20% were lifted to Wm82.v4 and Wm82.v6, respectively. The markers with MAFs lower than 1% were predominantly lifted at both Wm82.v4 and Wm82.v6, approximately 37%. Overall, the number of markers lifted to Wm82.v6 was slightly higher than that lifted to Wm82.v4 across other MAF ranges, particularly in the rare allele category, which showed MAFs below 1%. Next, we analyzed allele frequencies in the 497 PI soybean collection (Table 3).

**Table 3.** Categorization of lifted markers to 497 PI soybean collection using MAF.

| SNP Type | MAF | Number of SNPs |  |  |  |
| --- | --- | --- | --- | --- | --- |
|  |  | Wm82.v4 |  | Wm82.v6 |  |
| Lifted<br>to 497 PI<br>Soybean<br>Collection | ~0.01 | 31,407 | 18.89% | 31,447 | 18.89% |
|  | 0.01–0.05 | 36,235 | 21.79% | 36,245 | 21.78% |
|  | 0.05–0.10 | 23,918 | 14.38% | 23,952 | 14.39% |
|  | 0.10–0.20 | 31,230 | 18.78% | 31,264 | 18.78% |
|  | 0.20~ | 43,511 | 26.16% | 43,534 | 26.16% |
| **Total** |  | 166,301 | - | 166,442 | - |

Among the total lifted markers, 31,407 and 31,447 mapped to Wm82.v4 and Wm82.v6, respectively, corresponding to 18.89% of the dataset, were classified as rare alleles. Furthermore, 36,235 and 36,245 markers exhibited MAFs between 1% and 5%, 23,918 and 23,952 within 5%–10%, and 31,230 and 31,264 within 10%–20%. The number of markers with MAFs exceeding 20% totaled 43,511 and 43,534 for Wm82.v4 and Wm82.v6, respectively. Unlike the 430 Korean soybean core collection, where rare alleles were predominant, markers with MAFs greater than 20% comprised approximately 26% of the total in the PI collection, representing the most abundant category. Among the markers lifted to the 430 Korean soybean core collection and the 497 PI soybean collection, we also observed MAFs of markers that were uniquely lifted over to Wm82.v4 and Wm82.v6, respectively. The MAFs of rare alleles were higher in markers lifted over to Wm82.v6 in both collections (S5 and S6 Tables). Among the 380 uniquely lifted markers in Wm82.v6, 151 and 92 markers from the 430 Korean soybean core collection and 497 PI soybean core were classified as rare alleles, of which 86 and 52 were identified among the 239 uniquely lifted markers in Wm82.v4.

### Classification of lifted SNP array based on genomic features

Subsequently, we analyzed the positional characteristics of the SNP markers following liftover, focusing on their physical locations across chromosomes. The 175,202 and 175,763 markers lifted to Wm82.v4 and Wm82.v6, respectively, were predominantly located in the gene-rich chromosome arm regions (Fig 3).

**Fig 3.**
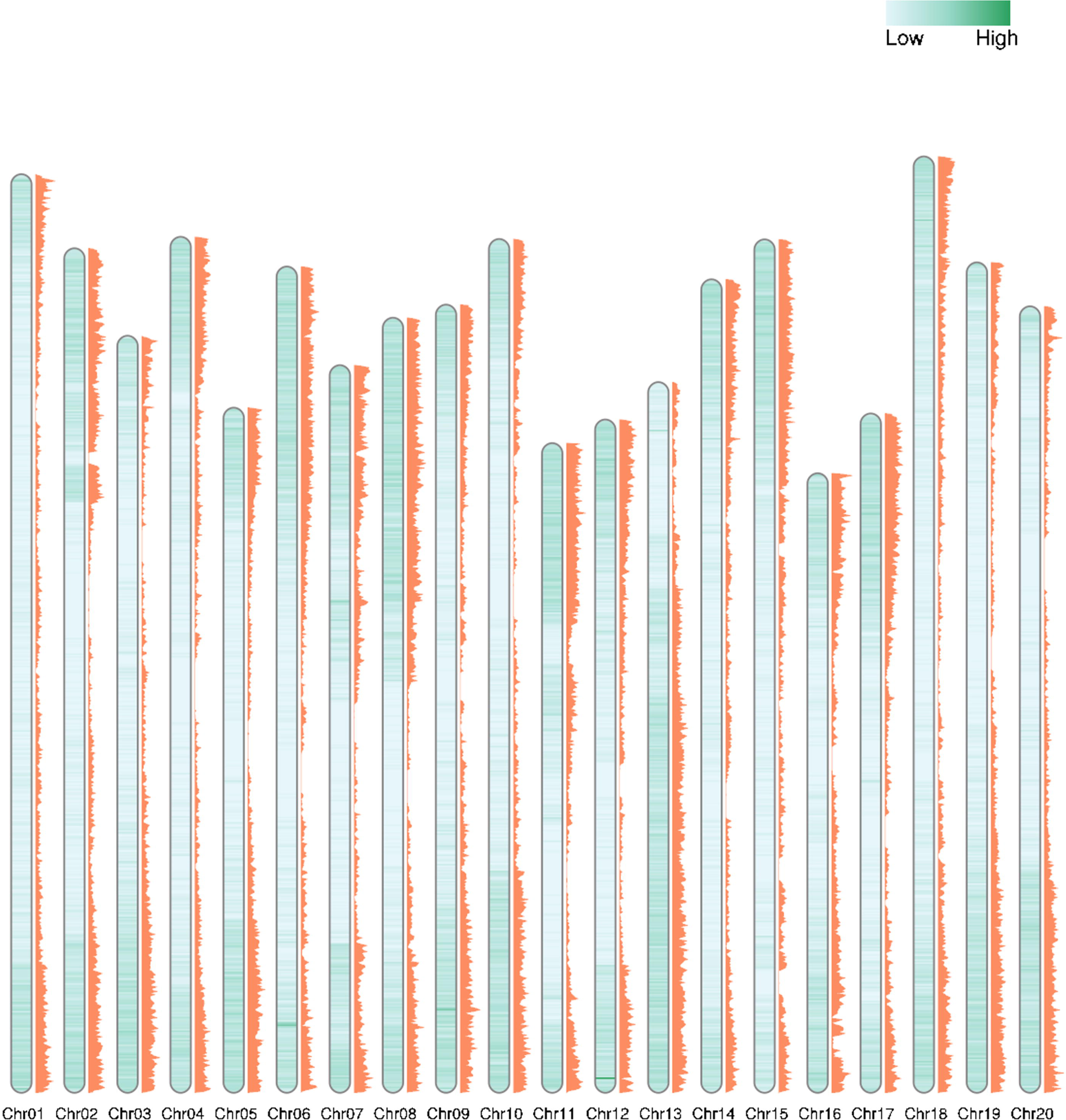
Comprehensive visualization of the gene distribution and lifted SNP marker landscape in the Wm82.v6. The inner green heatmap represents gene density along the chromosomes, while the adjacent orange histogram illustrates the density of lifted SNP markers.

For a more detailed classification, the lifted markers were categorized based on their chromosomal positions as follows: markers located within gene regions were classified as genic, those positioned outside gene regions but within 5 kbp upstream or downstream were classified as near-genic, and markers located in all other regions were classified as intergenic (Table 4). Markers lifted to Wm82.v4 and Wm82.v6 were predominantly located in genic regions. A total of 110,926 and 111,261 markers from Wm82.v4 and Wm82.v6, respectively, with approximately 60% of the total lifted markers, were located in genic regions. The next most common categories were intergenic and near-genic regions, with markers lifted to these regions in descending order of frequency. Among the three categories of lifted markers, the largest difference in liftover rates between Wm82.v4 and Wm82.v6 was observed in intergenic regions. In total, 37,492 and 39,066 markers were lifted to the intergenic regions of Wm82.v4 and Wm82.v6, representing 20.72% and 21.59%, respectively. Next, we identified the 5,759 and 5,198 markers that failed to be lifted to Wm82.v4 and Wm82.v6, respectively, which were already categorized as genic, intergenic, upstream, and downstream using Wm82.v1 from previous studies. Among the markers that failed to be lifted to both the Wm82.v4 and Wm82.v6 genome assemblies, the majority were predominantly located within the genic regions of Wm82.v1. A total of 3,555 and 3,329 markers were located in genic regions; 1,431 and 1,205 in intergenic regions; 263 and 219 in upstream regions; and 519 and 456 in downstream regions of Wm82.v1.

**Table 4.**
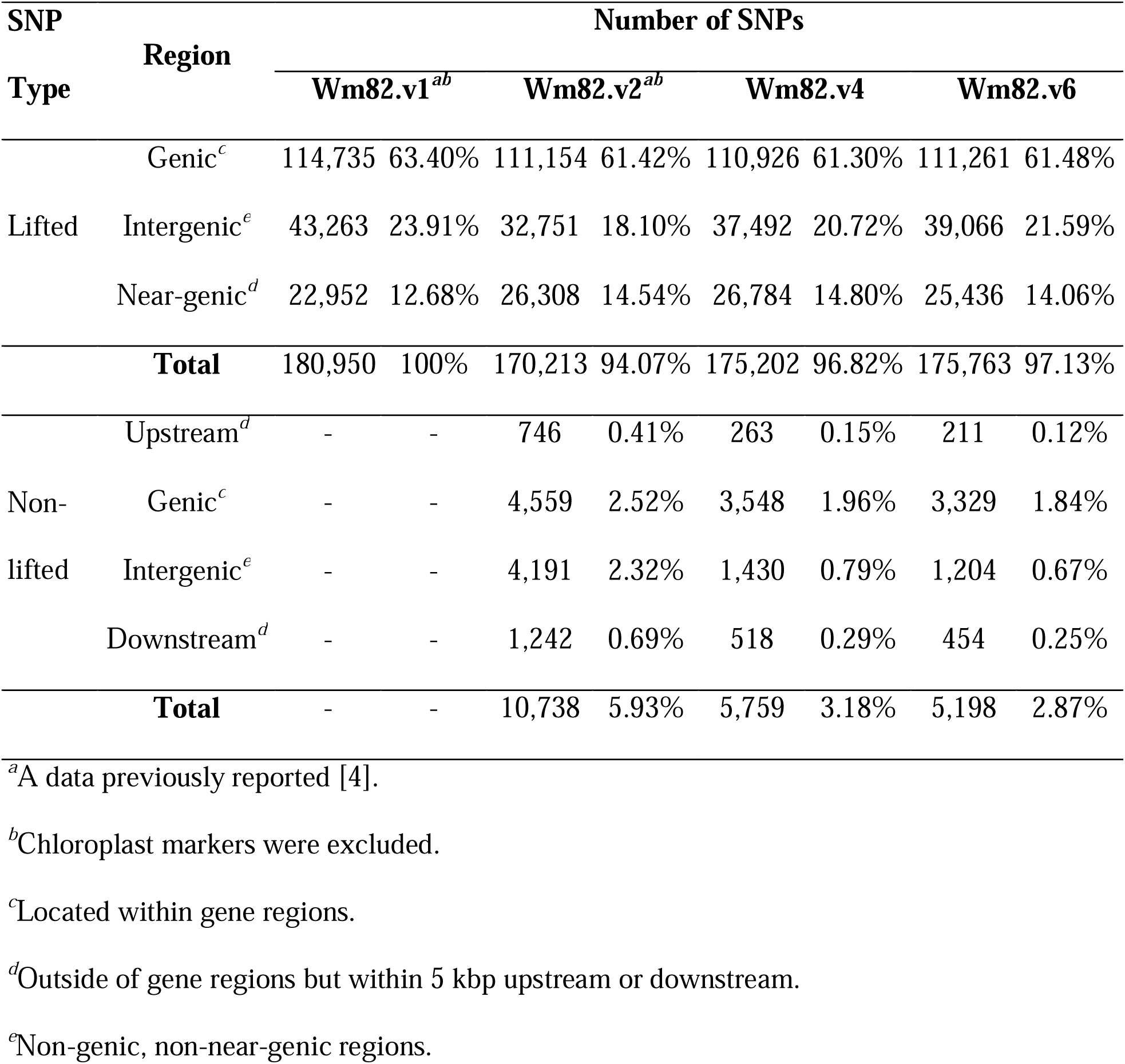
Distribution of the 180K SNP array markers at the various Wm82 genome versions.

| SNP Type | Region | Number of SNPs |  |  |  |  |  |  |  |
| --- | --- | --- | --- | --- | --- | --- | --- | --- | --- |
|  |  | Wm82.v1<sup>ab</sup> |  | Wm82.v2<sup>ab</sup> |  | Wm82.v4 |  | Wm82.v6 |  |
| Lifted | Genic<sup>c</sup> | 114,735 | 63.40% | 111,154 | 61.42% | 110,926 | 61.30% | 111,261 | 61.48% |
|  | Intergenic<sup>e</sup> | 43,263 | 23.91% | 32,751 | 18.10% | 37,492 | 20.72% | 39,066 | 21.59% |
|  | Near-genic<sup>d</sup> | 22,952 | 12.68% | 26,308 | 14.54% | 26,784 | 14.80% | 25,436 | 14.06% |
| **Total** |  | 180,950 | 100% | 170,213 | 94.07% | 175,202 | 96.82% | 175,763 | 97.13% |
| Non-lifted | Upstream<sup>d</sup> | - | - | 746 | 0.41% | 263 | 0.15% | 211 | 0.12% |
|  | Genic<sup>c</sup> | - | - | 4,559 | 2.52% | 3,548 | 1.96% | 3,329 | 1.84% |
|  | Intergenic<sup>e</sup> | - | - | 4,191 | 2.32% | 1,430 | 0.79% | 1,204 | 0.67% |
|  | Downstream<sup>d</sup> | - | - | 1,242 | 0.69% | 518 | 0.29% | 454 | 0.25% |
| **Total** |  | - | - | 10,738 | 5.93% | 5,759 | 3.18% | 5,198 | 2.87% |

<sup>a</sup>A data previously reported [4].
<sup>b</sup>Chloroplast markers were excluded.
<sup>c</sup>Located within gene regions.
<sup>d</sup>Outside of gene regions but within 5 kbp upstream or downstream.
<sup>e</sup>Non-genic, non-near-genic regions.

### Marker-lifted region comparison between the genomes

To compare Wm82.v4 and Wm82.v6, we visualized the genic SNP markers and their surrounding regions using their positions (Fig 4). Among the 863 markers that failed to be lifted to Wm82.v4, we focused on two markers: AX-90351012 and AX-90388399. These markers were previously classified as genic markers in Wm82.v1, and their associated genes were located in unplaced contigs in both Wm82.v1 and Wm82.v2. However, in Wm82.v6, these unplaced contigs and their corresponding markers were assigned to chromosome 10. Specifically, AX-90351012 and AX-90388399 were located within the gene *GmISU01.10G209200* in Wm82.v6. Synteny analysis revealed that the genes flanking *GmISU01.10G209200* were conserved between Wm82.v4 and Wm82.v6, although *GmISU01.10G209200* and *GmISU01.10G209300*, the genes nearest to *GmISU01.10G209200*, were present only in Wm82.v6.

**Fig 4.**
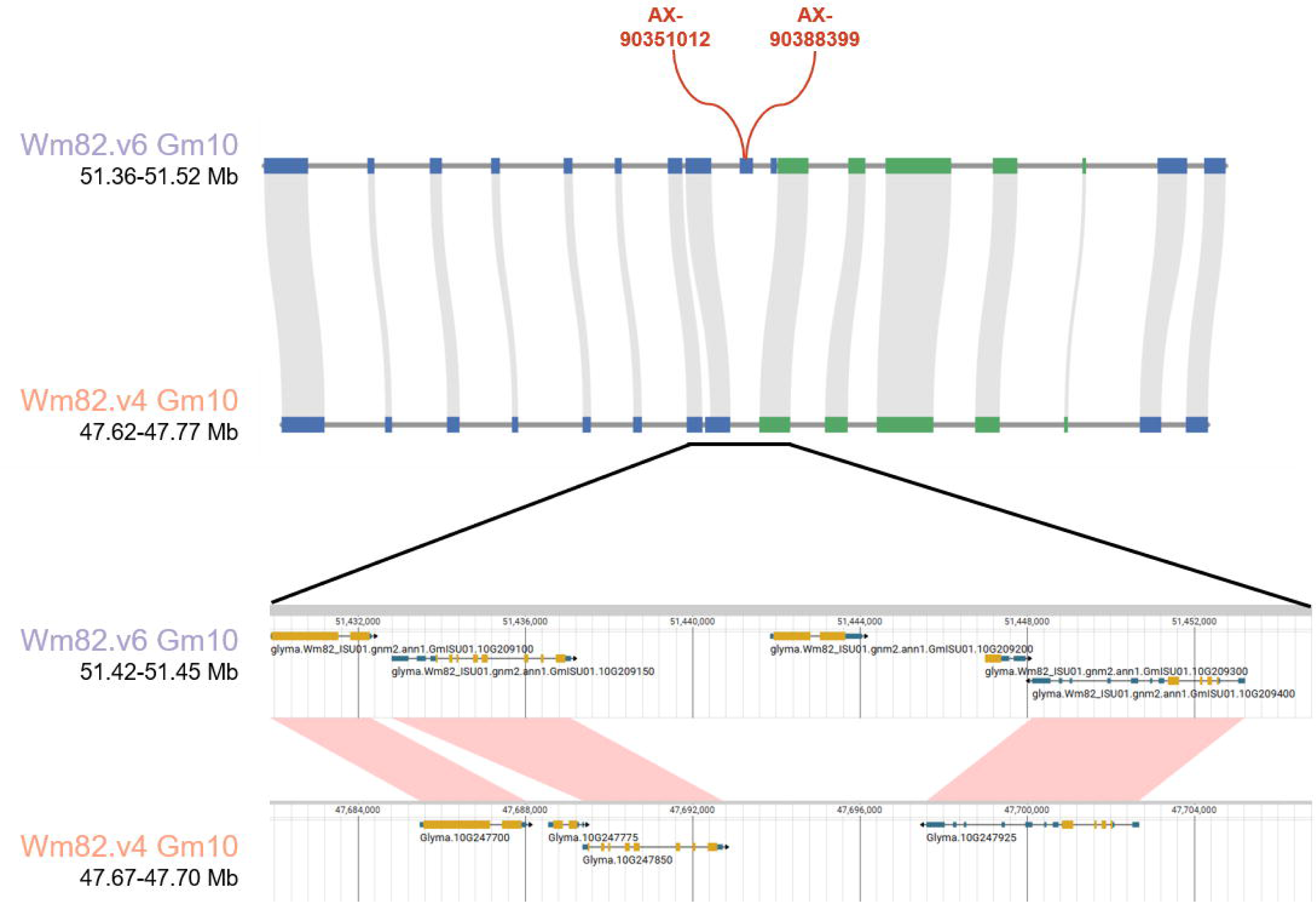
Visualization of genic regions for the liftover analysis comparing Wm82.v4 and Wm82.v6. Each block on the track represents the position of the gene. Gray and pink lines between tracks indicate syntenic regions, while the red line on the Wm82.v6 track marks the position of the genic marker.

To compare the intergenic regions between Wm82.v4 and Wm82.v6, we focused on the 3.85–4.45 and 4.00-4.60 Mbp regions on chromosome 3 from both genomes (Fig 5). In this region, 23 TEs longer than 2 kbp and 10 intergenic markers were identified in Wm82.v4, whereas 43 TEs longer than 2 kbp and 21 intergenic markers were identified in Wm82.v6. Among the 21 intergenic markers in Wm82.v6, 10 were located in regions homologous to Wm82.v4, whereas 11 were located in unique regions. Additionally, among the 21 intergenic markers of Wm82.v6 mapped to this region, six markers were nested within TEs. These marker-associated TEs were identified as belonging to the Gypsy retrotransposon superfamilies, as well as the Mutator and Helitron DNA transposon superfamilies. Of the six TE-nested markers, two were common between Wm82.v4 and Wm82.v6 and were located in homologous regions. The four remaining markers were uniquely mapped to Wm82.v6, which resides in unique regions.

**Fig 5.**
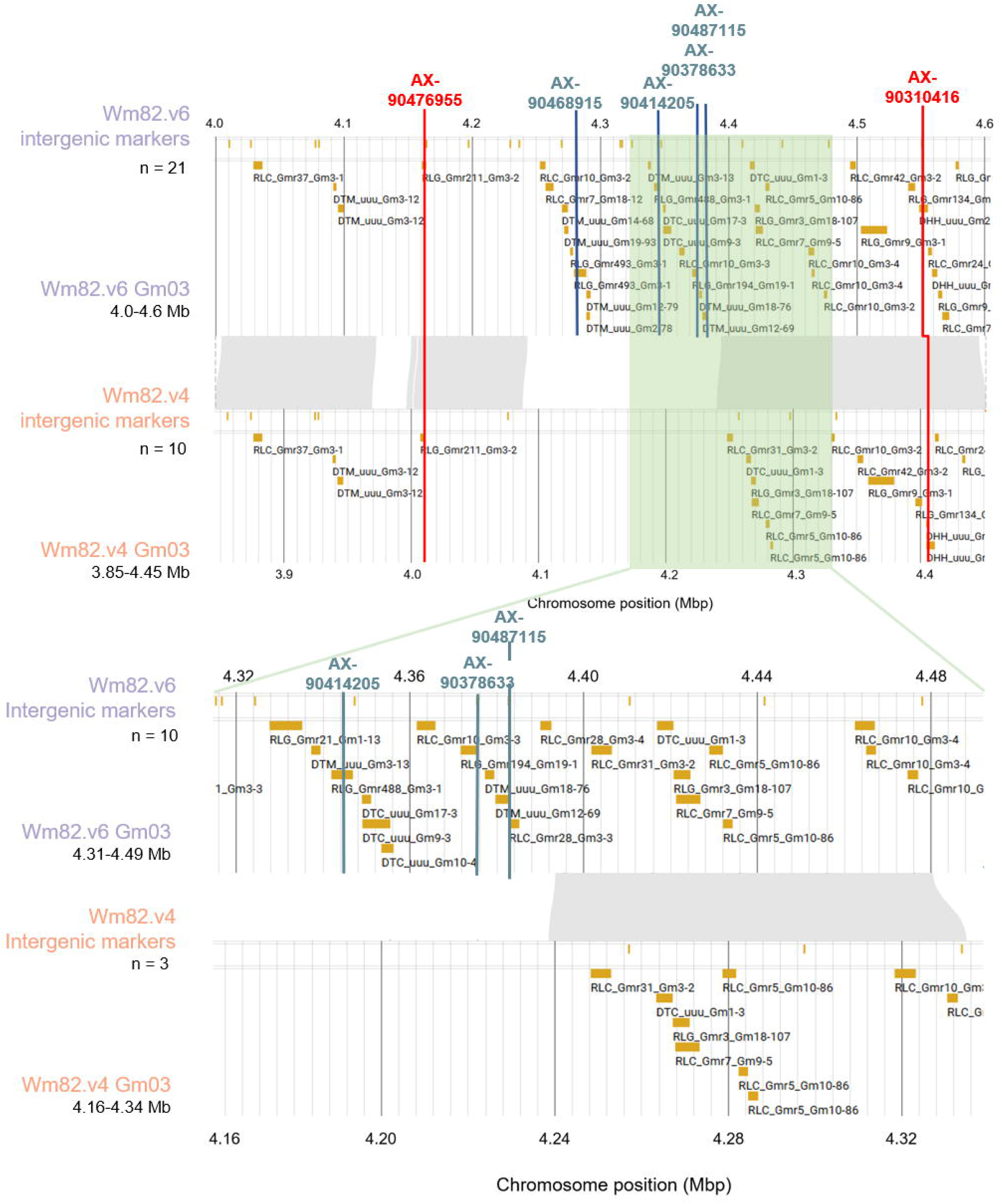
Visualization of intergenic marker regions for the liftover analysis comparing Wm82.v4 and Wm82.v6. The semi-transparent green blocks indicate zoomed-in regions. Gray regions between tracks indicate syntenic regions. The red lines on the track indicate TE nested intergenic markers specific to Wm82.v6, while the blue lines represent those present in both Wm82.v4 and Wm82.v6.

## Discussion

Williams 82, the soybean reference genome, was first assembled in 2010 and has since played a crucial role in advancing genomic studies within the soybean research community [5]. Early genotyping efforts, such as the 180K SoyaSNP project, relied on this genome for marker-based analyses [4]. However, the reference genome is continuously updated, and many of these markers remain anchored to early genome versions. This limits their utility in genomic studies. As genome assemblies have improved, particularly with the advent of T2T-level reference genomes, reassigning these markers to more complete assemblies has become increasingly important. In this study, we performed marker liftovers to Wm82.v4 and Wm82.v6, updated the marker positions to high-quality reference genomes, and emphasized the necessity of using the latest genome versions with the highest structural accuracy.

The liftover from Wm82.v1 to Wm82.v4 and Wm82.v6 revealed slightly higher success rates for Wm82.v6 on each chromosome. While most markers were retained on the same chromosomes across both versions, Wm82.v6 exhibited better liftover success, particularly for certain chromosomes. This improvement emphasizes the high accuracy and utility of Wm82.v6 for genomic analyses. However, a small proportion of the markers failed to be lifted to any version, possibly reflecting structural variations in the genome. These results underscore the need for updated reference genomes such as Wm82.v6 to ensure more precise and reliable genomic studies in soybean research.

The SNP markers lifted from the soybean collection to Wm82.v4 and Wm82.v6 further illustrate the value of updated reference genomes in capturing genetic diversity. Wm82.v6, in particular, showed a slightly higher number of lifted markers across all MAF ranges, with a more significant retention of markers in the rare allele category (MAF < 1%). This result suggests that advancements in genome assembly, such as those observed in Wm82.v6, enhance the detection of rare alleles, which is crucial for understanding genetic variation and selection. The improved retention of rare alleles in Wm82.v6 emphasizes the importance of using the most complete reference genome to improve marker-based analyses and population genetic studies.

The genomic distribution of the lifted SNP markers also highlights the broader impact of reference genome updates on marker positioning. In both Wm82.v4 and Wm82.v6, the majority of lifted markers were found in gene-rich chromosome arm regions, with approximately 60% located within genic regions. However, Wm82.v6 showed a higher proportion of markers retained in genic and intergenic regions than did Wm82.v4, suggesting enhanced annotation and assembly accuracy. The markers that failed to be lifted were predominantly located within the genic regions of Wm82.v1, potentially due to structural variations or assembly discrepancies. These findings emphasize the importance of using the most recent reference genome for accurate SNP mapping, particularly in functional genomics and marker-assisted selection studies.

Building on these observations, the comparison between Wm82.v4 and Wm82.v6 revealed notable improvements in marker liftover and genomic distribution. Markers such as AX-90351012 and AX-90388399, previously located on unplaced contigs in Wm82.v1, were successfully assigned to chromosome 10 in Wm82.v6. Synteny analysis showed that the genes flanking *GmISU01.10G209200* were conserved between the two versions, whereas the additional genes in Wm82.v6 highlighted its greater completeness than Wm82.v4. Furthermore, when comparing intergenic regions on chromosome 3, Wm82.v6 showed a higher number of TEs and intergenic markers than Wm82.v4, many of which were uniquely lifted or nested within TEs. Our results further underscore the superior genomic resolution and annotation accuracy of Wm82.v6, reinforcing the importance of using the latest reference genomes for more precise genetic analyses. By repositioning the markers based on the high-precision Wm82.v6 genome, we anticipate their usefulness in further genome-wide association studies and marker-assisted selection breeding efforts.

## Acknowledgments

This study was supported by the Faculty of Research Fund of Sejong University in 2024. This work was also carried out with the support of "Cooperative Research Program for Agriculture Science and Technology Development (Project No. RS-2023-00220176), Rural Development Administration, Republic of Korea.

## Supporting information

**S1 Table. Distribution of flanked marker sequence with identity below 100% S2 Table. Orientation of lifted markers relative to Wm82.v1**

**S3 Table. Chromosome-level assessment of the 180K SNP array data liftover efficiency relative to Wm82.v1**

**S4 Table. Liftover status of predicted markers from scaffolds in Wm82.v1 to chromosomes across different Wm82 genome versions**

**S5 Table. Categorization of uniquely lifted markers to 430 Korean soybean core collection using MAF**

**S6 Table. Categorization of uniquely lifted markers to 497 PI soybean collection using MAF**

## Data availability

The lifted Affymetrix Axiom® 180K SoyaSNP array data and genotype data of the Korean soybean core collection with 166,681 markers generated in this study are available at Figshare (<u>dx.doi.org/10.6084/m9.figshare.27991259</u>).

