## Supplementary Table 1-6 for "An updated resource of 180K soybean SNP genotyping array based on the T2T reference genome"

**Number of figures:** 0

**Number of tables:** 6

**S1 Table.**

| **Identity Range (%)** | **Wm82.v4** | **Wm82.v6** |
| --- | --- | --- |
| 70–90 | 0 | 2 |
| 50–70 | 275 | 280 |
| **Total** | 275 | 282 |

**S2 Table.**

| **Strand** | **Number of SNPs** | | |
| --- | --- | --- | --- |
|  | **Wm82.v2*^ab^*** | **Wm82.v4** | **Wm82.v6** |
| Forward | 159,641 | 162,020 | 162,758 |
| Reverse | 10,572 | 13,182 | 13,005 |
| **Total** | 170,213 | 175,202 | 175,763 |

*^a^*A data previously reported [4].

*^b^*Chloroplast markers were excluded.

**S3 Table.**

| **Chromosome** | **Total SNPs**  **(Wm82.v1)*^ab^*** | **SNPs lifted to the same chromosome in Wm82.v1** | | | | | |
| --- | --- | --- | --- | --- | --- | --- | --- |
|  |  | **Wm82.v2*^ab^*** | | **Wm82.v4** | | **Wm82.v6** | |
| Chr01 | 8,935 | 8,326 | 93.18% | 8,746 | 97.88% | 8,780 | 98.27% |
| Chr02 | 10,224 | 9,499 | 92.91% | 9,844 | 96.28% | 9,870 | 96.54% |
| Chr03 | 8,417 | 7,642 | 90.79% | 7,731 | 91.85% | 7,758 | 92.17% |
| Chr04 | 8,638 | 8,108 | 93.86% | 8,353 | 96.70% | 8,381 | 97.02% |
| Chr05 | 8,024 | 7,554 | 94.14% | 7,651 | 95.35% | 7,724 | 96.26% |
| Chr06 | 9,906 | 9,326 | 94.14% | 9,573 | 96.64% | 9,678 | 97.70% |
| Chr07 | 8,588 | 8,120 | 94.55% | 8,203 | 95.52% | 8,226 | 95.78% |
| Chr08 | 10,996 | 10,406 | 94.63% | 10,644 | 96.80% | 10,709 | 97.39% |
| Chr09 | 8,996 | 8,440 | 93.82% | 8,794 | 97.75% | 8,841 | 98.28% |
| Chr10 | 9,309 | 8,774 | 94.25% | 9,059 | 97.31% | 9,106 | 97.82% |
| Chr11 | 8,412 | 7,347 | 87.34% | 8,144 | 96.81% | 8,166 | 97.08% |
| Chr12 | 7,708 | 7,167 | 92.98% | 7,428 | 96.37% | 7,446 | 96.60% |
| Chr13 | 10,885 | 10,127 | 93.04% | 10,403 | 95.57% | 10,486 | 96.33% |
| Chr14 | 7,817 | 7,152 | 91.49% | 7,531 | 96.34% | 7,579 | 96.96% |
| Chr15 | 10,136 | 9,410 | 92.84% | 9,819 | 96.87% | 9,849 | 97.17% |
| Chr16 | 7,590 | 6,946 | 91.52% | 7,314 | 96.36% | 7,334 | 96.63% |
| Chr17 | 8,906 | 8,342 | 93.67% | 8,617 | 96.75% | 8,633 | 96.93% |
| Chr18 | 9,957 | 9,073 | 91.12% | 9,485 | 95.26% | 9,543 | 95.84% |
| Chr19 | 8,719 | 8,156 | 93.54% | 8,472 | 97.17% | 8,499 | 97.48% |
| Chr20 | 8,212 | 7,703 | 93.80% | 7,978 | 97.15% | 8,008 | 97.52% |
| **Total** | 180,375 | 167,618 | 92.93% | 173,789 | 96.35% | 174,616 | 96.81% |

*^a^*A data previously reported [4].

**S4 Table.**

| **Version** | **Wm82.v1*^a^****^b^* | **Wm82.v2*^a^****^b^* | | **Wm82.v4** | | **Wm82.v6** | |
| --- | --- | --- | --- | --- | --- | --- | --- |
| **Scaffold** | 575 | 142 | 24.70% | 51 | 8.87% | 0 | - |
| **Chromosome** | - | 313 | 54.43% | 350 | 60.87% | 456 | 79.30% |
| **Missing** | - | 120 | 20.87% | 174 | 30.26% | 119 | 20.70% |

*^a^*A data previously reported [4].

*^b^*Chloroplast markers were excluded.

**S5 Table.**

| **SNP Type** | **Minor Allele**  **Frequency** | **Number of SNPs** | | | |
| --- | --- | --- | --- | --- | --- |
|  |  | **Wm82.v4** | | **Wm82.v6** | |
| Lifted  to 430 Korean Soybean Core Collection | ~0.01 | 86 | 35.98% | 151 | 39.74% |
|  | 0.01–0.05 | 41 | 17.15% | 46 | 12.11% |
|  | 0.05–0.10 | 23 | 9.62% | 48 | 12.63% |
|  | 0.10–0.20 | 28 | 11.72% | 59 | 15.53% |
|  | 0.20~ | 61 | 25.52% | 76 | 20.00% |
|  | **Total** | 239 |  | 380 |  |

**S6 Table.**

| **SNP Type** | **Minor Allele**  **Frequency** | **Number of SNPs** | | | |
| --- | --- | --- | --- | --- | --- |
|  |  | **Wm82.v4** | | **Wm82.v6** | |
| Lifted  to 497 Plant Introduction Soybean Collection | ~0.01 | 52 | 21.76% | 92 | 24.21% |
|  | 0.01–0.05 | 55 | 23.10% | 65 | 17.11% |
|  | 0.05–0.10 | 27 | 11.30% | 61 | 16.05% |
|  | 0.10–0.20 | 41 | 17.15% | 75 | 19.74% |
|  | 0.20~ | 64 | 26.78% | 87 | 22.89% |
|  | **Total** | 239 |  | 380 |  |
